# Nanopore long-read only genome assembly of clinical Enterobacterales isolates is complete and accurate

**DOI:** 10.1101/2025.09.15.676237

**Authors:** Dorottya Nagy, Valentina Pennetta, Gillian Rodger, Katie Hopkins, Christopher R. Jones, The NEKSUS Consortium, Susan Hopkins, Derrick Crook, A. Sarah Walker, Julie Robotham, Katie L. Hopkins, Alice Ledda, David Williams, Russell Hope, Colin S. Brown, Nicole Stoesser, Samuel Lipworth

## Abstract

Whole bacterial genome sequence reconstruction using Oxford Nanopore Technologies (“Nanopore”) long-read only sequencing may offer a lower-cost, higher-throughput alternative for pathogen surveillance to ‘hybrid’ assembly with recent improvements in Nanopore sequencing accuracy. We evaluated the accuracy, including plasmid reconstruction, of Nanopore long-read only genome assemblies of Enterobacterales.

We sequenced 92 genomes from clinical Enterobacterales isolates, collected in England under a national surveillance program, with long-read Nanopore (R10.4.1, Dorado v5.0.0 super-high-accuracy basecalled) and short-read Illumina (NovaSeq) sequencing approaches. Genomes were assembled using three long-read only (Flye; Hybracter long; Autocycler), and three hybrid assemblers (Hybracter hybrid; Unicycler normal; bold). Three polishing modalities (Medaka v2 with subsampled or un-subsampled long-reads; Polypolish + Pypolca with short-reads) were investigated.

Autocycler circularised the most chromosomes (87/92 [95%]). Plasmid sequence reconstruction was comparable between all assemblers except Flye, all recovering 90-96% of plasmids, although the ‘ground truth’ was uncertain. Flye performed worse than other assemblers on almost all metrics. Autocycler + Medaka (un-subsampled long-reads) was the most accurate long-read only assembler/polisher combination, comparable to hybrid assemblies (median 0 [IQR:0-0] SNPs and 0 [IQR:0-1] indels per genome; quality value/Q score, 100 [IQR: 64-100]), with only 4/92 genome sequences having >10 SNPs/indels. Medaka polishing with un-subsampled long-reads resulted in small improvements in indels but not SNPs for both Flye and Autocycler assemblies. Seven-locus MLST, antimicrobial resistance, virulence, and stress gene annotation was equivalent across assembler/polisher combinations.

Nanopore long-read only bacterial genome assembly with Autocycler combined with Medaka polishing (using un-subsampled reads) is similarly accurate and possibly more complete than hybrid assemblies, representing a viable alternative for incorporating high-quality genomic data, including plasmids, into Enterobacterales surveillance.

**Data Summary:** Nanopore long-reads and Illumina short-reads from the 92 Enterobacterales isolates from this study have been uploaded to ENA (BioProject accession: PRJEB93885). Code for the Nextflow assembly pipeline, downstream analysis scripts, and R statistical analysis scripts are available on GitHub (https://github.com/oxfordmmm/NEKSUS_ont_hybrid_assembly_comparison). The following supplementary data tables are available on FigShare (https://figshare.com/account/home#/projects/253775):

- ENA Sample accessions and sample metadata (accessions_and_metadata.csv)
- Seqkit stats summaries of the Illumina and Nanopore reads (raw_qc_sup.cav)
- Summary of assembly contig features (contigs_summary_sup_cleaned.csv)
- Pairwise mash distances between contigs (mash_cleaned.csv)
- Plasmids matching across different assemblers compared to the Hybracter (hybrid) and manually-curated reference sets (plasmids_match_hybracter_mash.csv; plasmids_match_manual_mash.csv, respectively)
- Seven-locus multi-locus sequence type annotation (mlst_cleaned.csv)
- CheckM2 summaries of assemblies (checkm2_cleaned.csv)
- Nucleotide-level accuracy of assemblies (SNP, Indels, and Quality value compared to short-read mapping; assembly_nucleotide_accuracy_cleaned.csv)
- Bakta annotation (bakta_by_contig_cleaned.csv)
- AMRFinderPlus annotations of contigs (amrfinder_plus_cleaned.csv)
- MOB-suite annotation summaries of contigs (mobsuite_cleaned.csv)

**Impact Statement:** Nanopore long-reads have historically been too error-prone to use alone for accurate bacterial genome assembly, necessitating additional Illumina short-reads to achieve structurally complete and accurate ‘hybrid’ genome assemblies for public health surveillance. This increases cost and complexity. Previous studies have shown that recent improvements in Nanopore chemistry (R10.4.1 flowcell) and basecalling (super-high accuracy) allow high-quality long-read only assemblies on a small number of laboratory reference strains. This is the first evaluation, to our knowledge, to assess Nanopore long-read only genome assembly compared with hybrid assembly on a large number of clinical isolates. In addition, this is the first large-scale evaluation of the recently released automated consensus long-read assembly tool, Autocycler.

We show that Autocycler long-read only assemblies are more structurally complete for chromosomal sequences, while reconstructing a similar number of plasmids to other long-read and hybrid assemblers. Most long-read polished, Autocycler-assembled genome sequences have 0 errors (median: 0 SNPs/indels) relative to a short-read polished (hybrid) Autocycler assemblies, enabling accurate annotation of key genes.

## Introduction

Hybrid assembly combining short- and long-read genomic sequencing is widely used in research to assemble complete and accurate bacterial genome sequences. Incremental improvements in Nanopore flowcells/chemistry (10.4.1 flowcell/kit 14) and basecalling accuracy (Dorado v5.0.0 super-high accuracy DNA model)(1–5) have been shown in small-scale evaluations to facilitate long-read only assemblies that may now be comparable in accuracy to hybrid assembly(6, 7). Nanopore-only sequencing may also offer advantages over hybrid sequencing, including cost effectiveness, real-time data generation and decentralised implementation(8, 9).

Highly accurate bacterial genome reconstruction, with minimal noise from sequencing artefact, is key for identifying closely-related clusters of isolates and plasmids for outbreak detection(10). Accurate reconstruction of mobile genetic elements (MGEs) such as plasmids in particular, is clinically and epidemiologically important as plasmids are common transmission vectors for antimicrobial resistance (AMR) genes in clinically-relevant Enterobacterales(11, 12). Long-read or hybrid assembly approaches can facilitate plasmid sequence reconstruction and therefore analysis of AMR gene epidemiology compared to short-reads, which may not be able to resolve highly repetitive sequences often associated with MGEs(13, 14). Nevertheless, Nanopore-only genome assembly accuracy has only been validated for a small number of reference bacterial isolates(15, 16), and has not yet been assessed on a large collection of clinical isolates, including for plasmids as well as chromosomes. This may be important because of the reliance of long-read basecalling models on training datasets of unknown size and diversity, whose performance may therefore generalise poorly to clades not included in these training datasets. Similarly, although best-practice assembly guidelines have been proposed(6, 17, 18), multiple long-read assembly pipelines implement these guidelines with slight variations(16, 19–22), and no robust consensus exists, particularly regarding the optimal strategy for plasmid assembly.

In this study, we comprehensively evaluated the completeness and accuracy of 92 Nanopore long-read only assemblies (with and without polishing) compared to hybrid assembly in reconstructing both chromosomes and plasmids using isolates collected in The <u>N</u>ational *<u>E</u>scherichia coli* and *<u>K</u>leb<u>S</u>iella* spp. bloodstream infection (BSI) and Carbapenemase-producing Enterobacterales (CPE) <u>U</u>K <u>S</u>urveillance (NEKSUS) study.

## Methods

### Isolate collection

Nine English NHS Trusts (groups of hospitals under the same administration) representing the largest in terms of number of emergency admissions across all seven NHS England regions were recruited to the NEKSUS consortium. Consecutive, unselected BSI and CPE-positive rectal screening isolates were collected between October 2023 and March 2024 as part of routine clinical practice. One convenience sample of the first 96 Enterobacterales isolates collected, mostly *E. coli* and *Klebsiella* spp. (Table S1), sequenced from three regions, were included in this analysis as our isolates were sequenced in batches of 96. Isolates were stored in brain-heart infusion (BHI) broth with 10% glycerol at -70°C, then grown on blood agar for 24h at 37°C, following which a colony sweep of the pure bacterial culture was suspended in 1 ml phosphate buffer saline, pelleted, and cold-packed. Bacteria were subcultured for a further 24h at 37°C where there was insufficient growth after 24h.

### DNA extraction and sequencing

DNA extraction, library preparation and sequencing were conducted at GENEWIZ Germany GmbH (Leipzig, Germany). DNA was extracted using the MagMAX Microbiome Ultra Nucleic Acid Isolation Kit with bead plate (Life Technologies, Carlsbad, CA, USA). Genomic DNA was quantified using the Qubit 4.0 Fluorometer and qualified using the Agilent 5600 Fragment Analyzer. The same DNA extract was sequenced by both methods.

For Nanopore sequencing the Rapid Barcoding Kit 96 V14 (Oxford Nanopore Technologies, Oxford, UK) was used according to the manufacturer’s recommendations. Briefly, sequencing libraries were generated using a transposase, which simultaneously cleaves template molecules and attaches barcoded tags to the cleaved ends. The barcoded samples were then pooled (96-plexed) before solid phase reversible immobilisaton (SPRI)-cleaning and addition of Rapid Adapters to the tagged ends. The library pools were loaded onto ONT PromethION flow cells (R10 [M Version]) – one 96-plex pool per flow cell – and sequenced on a PromethION P2 Solo for 72 hours according to the manufacturer’s instructions.

For Illumina sequencing the NEBNext Ultra II DNA Library Prep Kit for Illumina (New England Biolabs, Ipswich, MA, USA), including clustering and sequencing reagents, was used according to manufacturer’s recommendations. Briefly, the genomic DNA was fragmented by acoustic shearing with a Covaris LE220 instrument. Fragmented DNA was cleaned up and end repaired. Adapters were ligated after adenylation of the 3’ ends followed by enrichment by limited cycle PCR. DNA libraries were validated using the Agilent TapeStation (Agilent Technologies, Palo Alto, CA, USA), and were quantified using a Qubit 4.0 Fluorometer. The libraries were multiplexed on a flowcell and loaded on the Illumina NovaSeq X Plus instrument according to manufacturer’s instructions. The samples were sequenced using a 2×150bp paired-end (PE) configuration. Raw sequencing data (.bcl files) generated from Illumina NovaSeq were converted into fastq files and de-multiplexed using Illumina’s bcl2fastq(23) v2.20 software.

### Bioinformatic analysis

Computational analysis was performed on a virtual machine in the Oracle Cloud Infrastructure. POD5 files were basecalled and demultiplexed using Dorado(24) v5.0.0 (super high accuracy 5mCG, 5hmCG and 6mA methylation aware simplex DNA model). All bioinformatic tools were run using default settings unless otherwise specified. Raw-read quality was evaluated with SeqKit(25) v2.9.0. Long-reads were randomly subsampled to 60x using the built-in subsampling and genome size estimation scripts from Autocycler(20) v0.2.1, and short-reads were randomly subsampled to 100x (50x for each paired-end read) with Rasusa(26) v2.1.0. Genome sequences were assembled using three long-read only assemblers (Flye(27) v2.9.5, Hybracter(19) (long) v0.11.2, the consensus assembler Autocycler(20) v0.2.1), and three hybrid assemblers (Hybracter(19) (hybrid), Unicycler(7) v0.5.1 (normal and bold modes)). The input long-read assemblies used for Autocycler were four assemblies each of Canu(28) v2.2, Flye(27), Raven(29) v1.8.3, Miniasm(30) v0.3, and Hybracter(19) (long) (which incorporates the plasmid assembly tool Plassembler(31)), where each of the four assemblies was derived from a randomly subsampled set of reads. The Flye and Hybracter (long) assemblies from the first subsampled read set were used in downstream analyses. Three polishing modalities were investigated: long-read polishing with one round of Medaka(32) v2.0.1 using 1) subsampled long-reads, 2) un-subsampled long-reads, or 3) short-read polishing with Polypolish(33) v0.6.0 and Pypolca(17, 34) v0.3.1 (‘--careful’ flag; Fig. 1).

**Figure 1:**
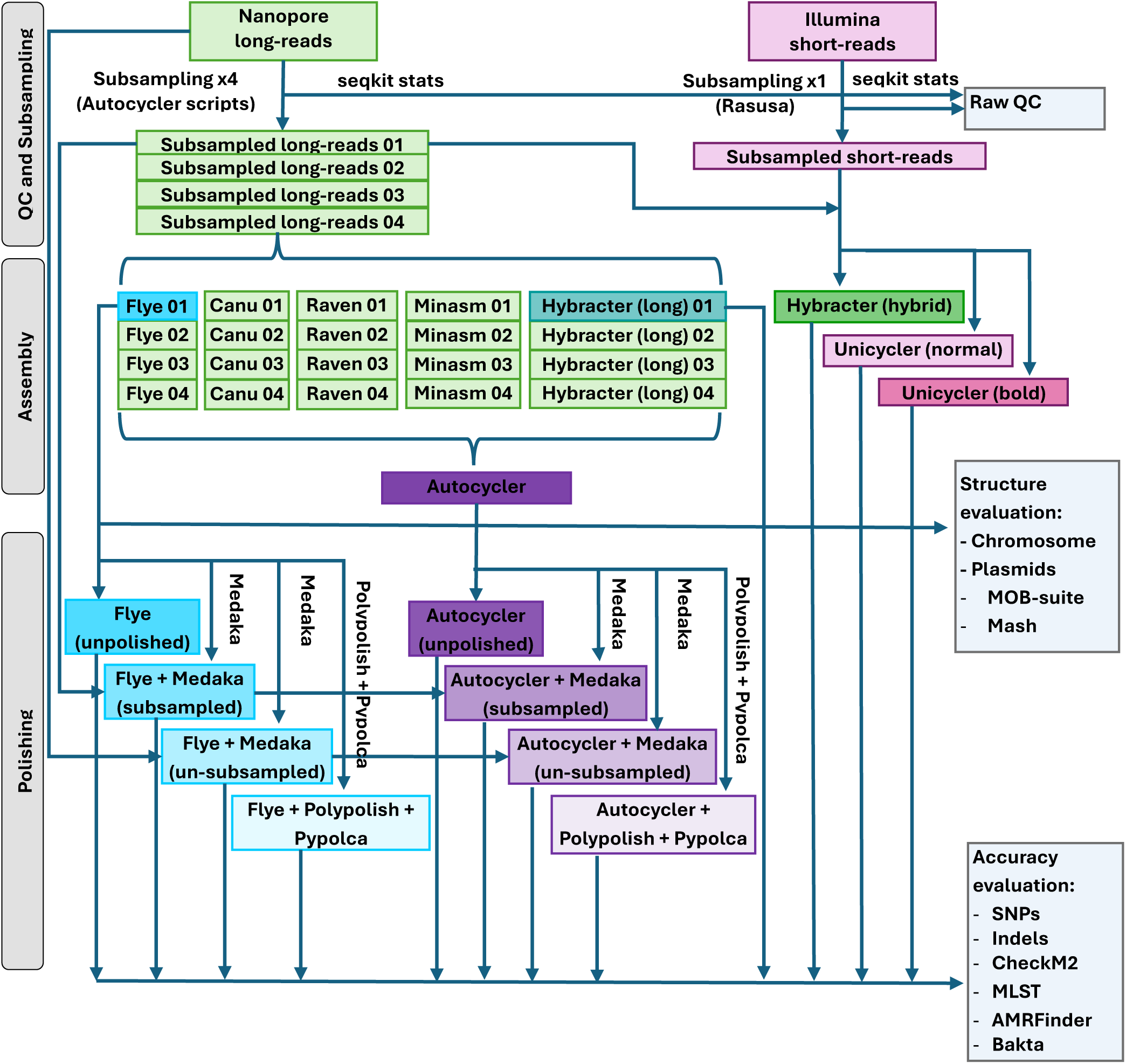
Schematic diagram of assembly, polishing and downstream analysis pipeline.

### Assembly quality control

Quality control of assemblies was done using SeqKit(25) stats and CheckM2(35–37) v1.0.2, excluding isolates where any assembly for that isolate had <99% completeness and/or >5% contamination. 4/96 (4%) isolates had >5% ‘contamination’ based on the checkM2 output, likely corresponding to mixed isolate sequences (i.e. not pure cultures), so were excluded from subsequent analyses. The remaining 92/96 (96%) pure-culture isolates passed the completeness threshold.

### Assembly annotation

Assemblies from all 12 assembler/polisher combinations were annotated using Bakta(38) v1.10.4, 7-locus MLST (mlst(39) v2.23.0), AMRFinderPlus v4.0.3 (species flag inferred from Kraken2(40) v2.1.3) and MOB-suite(41, 42) v3.1.9 (mob_recon and mob_typer).

### Chromosome evaluation

Assemblies from the six different assemblers (without polishing) were evaluated for structural completeness of chromosomes and plasmids, as polishing is not expected to alter structure. Chromosomes were considered ‘fully reconstructed’ if the chromosomal contig was >4Mb and circularised.

### Plasmid evaluation

Contigs ≥1,000bp and ≤400,000bp in length were considered potential plasmids. Mash distances between all potential pairwise plasmid combinations were calculated using Mash(43, 44) v2.3 (k-mer size = 21, sketch size 10,000,000).

Plasmid reconstruction was assessed by comparing with two alternative ‘reference’ plasmid sets generated from the assembly data in this study, due to the absence of a ‘ground truth’ for these isolates. The first ‘reference’ plasmid set included all circular potential plasmids recovered by Hybracter (hybrid), which incorporates the plasmid assembly tool Plassembler(31), recommended in best-practice assembly guidance(6). The second ‘reference’ plasmid set was created using a manually-curated consensus approach considering all six assemblies for each isolate. This latter manually-curated reference set was constructed by matching each potential plasmid contig from the six assembly methods to its most similar contig from each other assembler based on mash distance, forming a network with all pairwise assembler combinations. The R package igraph(45, 46) v2.1.4 was used to extract connected components (sub-networks within each sample with at least one mash-distance connection between nodes). Each connected component was assigned a ‘match-set’ ID. Three (out of 303) match-sets (connected components) contained more than one contig per assembler, and were corrected manually (two were likely partial plasmids and one was likely a chimeric Unicycler (bold) plasmid that joined two separate plasmid match-sets together; data not shown). ‘True’ match sets were retained in the manually-curated reference set where at least two assemblers’ contigs were present, circular, of similar length (±10%) and had a low mash distance (<0.025). The 0.025 mash distance threshold reflects the highest possible mash distance between draft and complete plasmid assemblies of the same plasmid from the original MOB-suite publication(42).

Plasmid reconstruction for each assembler was then evaluated, for the Hybracter (hybrid) reference set, by matching potential plasmid contigs to each reference plasmid set based on circularity (i.e. circular or linear), length (±10%), and mash distance (<0.025). Plasmids were ‘present’ if all three match criteria were met, or ‘misassembled’ if at least one of the criteria were not met. Plasmids were ‘absent’ if none of the criteria were met, if only the circularity matched (but not length or mash distance), or if no contig from an assembler could be matched to that set. For the manually-curated reference set, where no single reference plasmid was available, mash distance and length similarity criteria were fulfilled if an assembler’s plasmid matched more than half of the other plasmids in a match set (see supplementary data file plasmids_mash_manual_mash.csv).

### Nucleotide-level accuracy

Nucleotide-level accuracy was assessed in a reference-free manner by aligning Illumina short-reads to the 12 assembler-polisher combinations using the Pypolca(17) in-built read aligner and variant caller (BWA(47) 0.7.18 and Freebayes(48) v1.3.6). Single nucleotide substitutions (SNPs), short insertions/deletions (indels) and quality value (QV) were extracted from the .vcf output file. QV, like Phred score, is a measure of accuracy where higher QV signifies a more accurate consensus (QV = -10 * log10(probability of error), where a 0-error probability takes the value of Q100). Mean gene length was extracted from CheckM2(37) as a further measure of accuracy. Errors may introduce premature stop codons and are thus expected to reduce the length of coding sequences(38).

### Statistical analyses and visualisations

Statistical analysis and visualisation were done in R(49) v4.4.1 using ggplot2(50) v3.5.1 and other tidyverse(51) v1.3.1 functions, gridExtra(52) v2.3, cowplot(53) v1.1.3, psych(54) v2.5.6 and irr(55) v0.84.1 packages. Global test for uneven proportions in categorical variables was done using the multiple-group Fleiss’ Kappa test, and for continuous variables, with a Friedman test to account for non-independence between different assemblers’ ‘observations’ on the same isolate. Pairwise test between assemblers for differences in proportions were done using McNemar’s Χ^2^-test with continuity correction and for differences in counts, with Wilcoxon signed-rank tests. A Bonferroni correction was applied to all pairwise testing to account for multiple testing. An exact binomial test was used to test for a significant difference to 1 for the proportion of plasmids reconstructed compared to the Hybracter (hybrid) reference set. Clinker(56) v0.0.31 was used to visualise plasmid alignments using the Bakta(38) v1.10.4 annotated .gbff files.

## Results

### Raw sequences

#### High sequencing depth and quality was achieved for both Illumina short- and Nanopore long-reads

Over 200x sequencing depth was achieved for both Illumina and Nanopore reads (Table S2). Median long-read length was 5814bp (IQR: 5366-6338), and median estimated Phred quality score was 16.6 (IQR: 16.4-16.8). Subsampling did not affect median read length or read quality (Table S2; Fig. S1).

### Structural completeness

#### Chromosome reconstruction was optimal using the consensus long-read only assembler, Autocycler

Autocycler circularised the most chromosomal sequences, 95% (87/92), significantly more than Unicycler (80% [74/92], pairwise McNemar’s *p=0.006*), Unicycler bold (85% [78/92], *p=0.039*) and Flye (85% [78/92], *p=0.027*), Hybracter (hybrid) (86% [79/92], *p=0.043*), while there was no statistical evidence of a difference to Hybracter (long) (87% [80/92], *p=0.070;* Table 1; Fig. 2a). Notably, for two isolates that were correctly assembled by all other assemblers, Autocycler failed to generate a circular consensus chromosome (Fig. 2a), producing highly fragmented draft assemblies instead.

**Figure 2:**
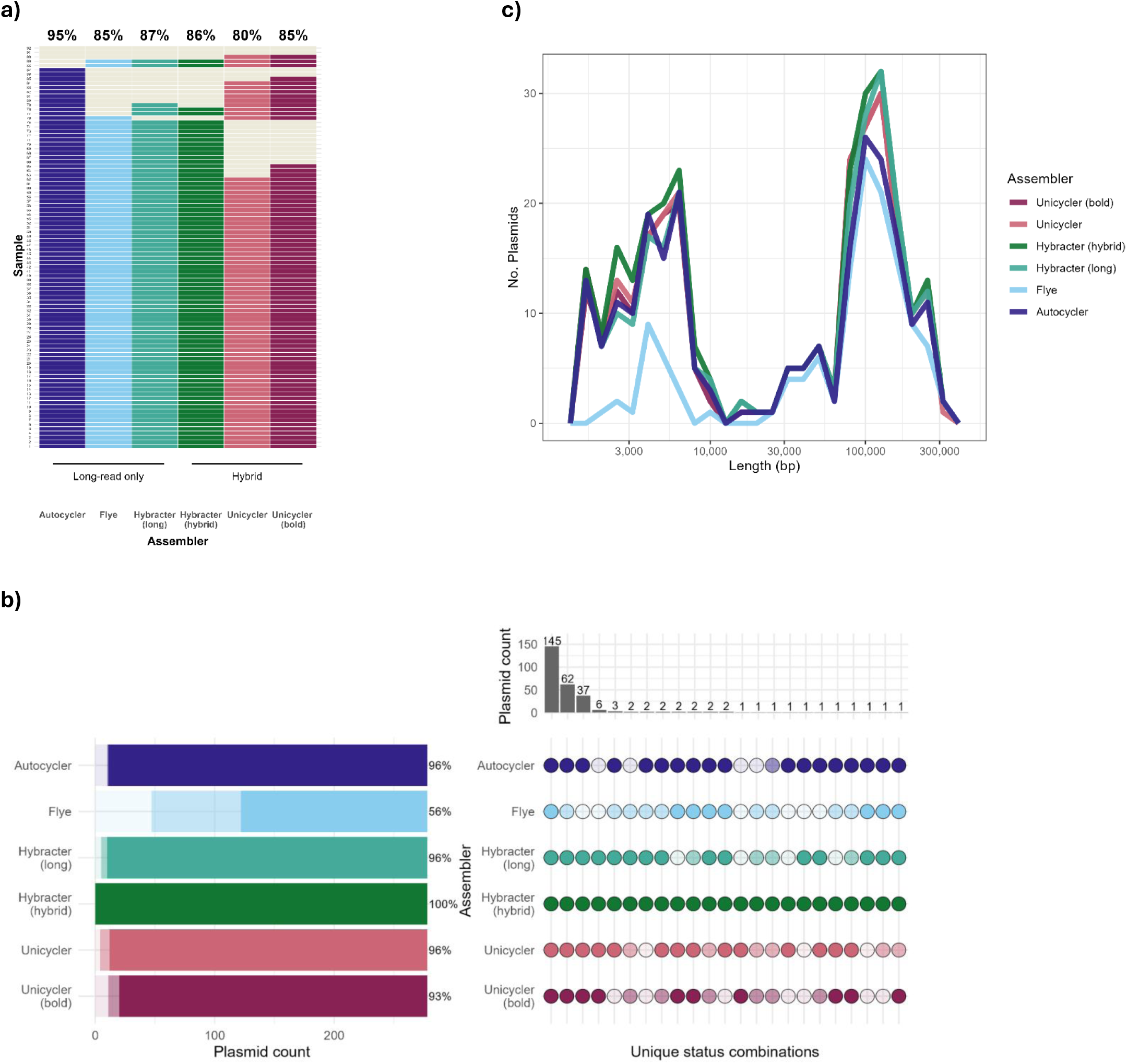
Structural completeness of 92 pure culture Enterobacterales genome sequences assembled by different long-read only and hybrid assemblers. Genome sequences were assembled using Dorado v5.0.0 super-high accuracy basecalled Nanopore long-reads, plus Illumina short-reads for hybrid assembly. **a)** Number and percentage of isolates with a fully circularised chromosome (dark-coloured tiles) or an incompletely circularised chromosome (light cream tiles) by assembler. **b)** Upset plot of plasmid assembly status combinations across assemblers. Plasmid sequence reconstruction (assembly status) is compared to a Hybracter (hybrid) plasmid reference dataset, defined as circular contigs ≤400,000bp and ≥1,000bp assembled by Hybracter (hybrid)(n=278) across the 92 Enterobacterales isolates analysed. Dark circles represent ‘present’ plasmids where length (±10%), mash distance (<0.025) and circularity all matched the Hybracter (hybrid) ‘reference’ plasmid, lighter colours indicate misassembled plasmids, where the length difference was >10%, mash distance >0.025, or the contig was non-circular and the palest shades indicate absent plasmids, where no contig was found matching other plasmids in the reference plasmid set. **c)** Frequency polygon of length distribution of ‘present’ plasmids by assembler.

**Table 1:** Chromosomal sequence circularisation and accuracy of plasmid sequence reconstruction for different assemblers using Dorado v5.0.0 super-high accuracy basecalled Nanopore long-reads. Plasmid sequence reconstruction was compared with the Hybracter (hybrid) plasmid reference dataset, defined as circular contigs ≤400,000bp and ≥1,000bp assembled by Hybracter (hybrid)(n=278) across 92 Enterobacterales isolates analysed; the denominator for plasmids was therefore 278 throughout.

|  | Assembler |  |  |  |  |  | p-value<sup>†</sup> |
| --- | --- | --- | --- | --- | --- | --- | --- |
|  | Autocycler<br>n (%) | Flye<br>n (%) | Hybracter<br>(long)<br>n (%) | Hybracter<br>(hybrid)<br>n (%) | Unicycler<br>n (%) | Unicycler<br>(bold)<br>n (%) |  |
| **Chromosomes circularised (N=92)** | 87 (94.6%) | 78 (84.8%) | 80 (87.0%) | 79 (85.9%) | 74 (80.4%) | 78 (84.8%) | <0.0001 |
| **Present\* plasmids (N=278)** | 267<br>(96%) | 156<br>(56.1%) | 268<br>(96.4%) | 278<br>(100%) | 266<br>(95.7%) | 258<br>(92.8%) | 0.002 |
| **Misassembled\*\* plasmids (N=278)** |  |  |  |  |  |  |  |
| Non-circular | 0 (0%) | 16 (5.8%) | 2 (0.7%) | 0 (0%) | 4 (1.4%) | 2 (0.7%) |  |
| Length mismatch | 1 (0.4%) | 41 (14.7%) | 1 (0.4%) | 0 (0%) | 1 (0.4%) | 1 (0.4%) |  |
| Mash distance >0.025 | 0 (0%) | 1 (0.4%) | 0 (0%) | 0 (0%) | 0 (0%) | 0 (0%) |  |
| Non-circular & length mismatch | 0 (0%) | 17 (6.1%) | 2 (0.7%) | 0 (0%) | 3 (1.1%) | 6 (2.2%) |  |
| **Absent plasmids (N=278)** | 10 (3.6%) | 47 (16.9%) | 5 (1.8%) | 0 (0%) | 4 (1.4%) | 11 (4%) |  |

\*'Present' plasmids are defined as contigs meeting all three match criteria: circular, length within 10% and mash distance <0.025 of a Hybracter hybrid reference plasmid.
\*\*Misassembled plasmids are defined as contigs that failed to meet at least one of the matching criteria, or were non-circular and a different length (>10% difference).
<sup>†</sup>p-value for Fleiss' Kappa test for uneven proportions of circularised chromosomes or 'present' plasmids across all assemblers.

#### Plasmid reconstruction was improved by Autocycler or Hybracter compared with Flye

Given the absence of a ‘ground truth’ for plasmids in the sequenced isolates, we considered two ‘reference’ plasmid sets generated from the assembly data. The first was the Hybracter (hybrid) reference set, and the second, a manually-curated reference set considering potential plasmids across all assemblers. All plasmids from the Hybracter (hybrid) reference set (n=278) were present in the manually-curated set. However, the manually-curated set included an additional 25 plasmids (total 303 vs 278 plasmids), which were missing from the Hybracter (hybrid) reference set, mostly due to being non-circular (17/25, 68%), or non-circular and of different length (3/25, 12%), while 5/25 (20%) plasmid sets could not be matched to any Hybracter (hybrid) contigs not already in another match set (all pairwise mash distances >0.2; Table S3).

Compared with the Hybracter (hybrid) reference set, Flye reconstructed significantly fewer plasmids than all the other assemblers (56% [156/278]; exact binomial test *p<0.0001* vs 100% reconstructed by Hybracter (hybrid) and McNemar’s *p<0.0001* vs Autocycler, Hybracter (long), Unicycler, and Unicycler (bold)). Among the remining assemblers, 93-96% of plasmids were reconstructed, which was significantly fewer than 100% of the Hybracter (hybrid) reference set (all exact binomial test *p<0.0001*; Table 1; Fig. 2b). There was no evidence of a difference between the 96% (267/278) of plasmids reconstructed by Autocycler compared to the other assemblers besides Flye (Hybracter (long) 96% [268/278], McNemar’s *p=1* vs Autocycler, Unicycler 96% [266/278], *p=1* and Unicycler (bold) 93% [258/278], *p=0.095*).

Similarly, compared with the manually-curated reference set, Flye reconstructed significantly fewer plasmids than all other assemblers (55% [166/303]; pairwise McNemar’s *p<0.0001* vs each of the five other assemblers). Flye more frequently missed or misassembled small, <10,000bp, plasmids (Fig. 2c; S2b), and incorrect length was the most common reason for Flye plasmid misassembly (Table 1; S2). Among the remaining assemblers, 90-94% of plasmids were reconstructed compared to the manually-curated reference set. Autocycler reconstructed 94% (285/303) of plasmids, significantly more than Hybracter (long) (90% [272/303]; McNemar’s *p=0.014*). However, there was no evidence of a difference between the number of plasmids reconstructed by Autocycler compared to the other assemblers: Hybracter (hybrid) (91% [276/303]; McNemar’s *p=0.066* vs Autocycler), Unicycler (93% [282/303]; *p=1*), or Unicycler (bold) (90% [274/303]; *p=0.296*; Table S3; Fig. S2a).

Of the 10 Autocycler plasmids with a mash distance of 0 to the corresponding Hybracter (hybrid) plasmid, 2/10 had a missing MOB-suite IncFIC replicon annotation despite identical sequence (Fig. S3). In both cases, the Autocycler plasmid was reversed (i.e. the reverse complement strand was represented in the fasta file) compared with the other plasmids. The Flye plasmid sequence was also missing an IncFIC annotation in one of these two plasmids; however, this difference was not observed in the other 232 plasmids across other assemblers with a mash distance of 0 to the Hybracter (hybrid) reference.

### Assembly accuracy

#### Unpolished Autocycler assemblies are more accurate than non-consensus long-read assemblers, while differences compared with hybrid assemblers are small

Autocycler was the most accurate long-read only assembler, with 37% of unpolished assemblies (34/92) having 0 SNPs or indels when compared with 11% (10/92) for unpolished Flye and 7% (6/92) for Hybracter (long). For unpolished Autocycler, this equated to a median of 0 SNPs/Mb (IQR: 0-0.17) and 0.18 indels/Mb (IQR:0-0.39), and a median quality value (QV) of Q67 (IQR:63-100; Fig. 3a-c; Table S4). The differences in accuracy between unpolished Autocycler, unpolished Flye or Hybracter (long) were significant (pairwise Wilcoxon signed rank *p<0.0001* for SNPs, indels and QV), while there was no evidence of a difference in accuracy between unpolished Autocycler and Unicycler (normal or bold mode; *p=1* for all metrics). There was no evidence of a difference between Flye and Hybracter (long) assemblies (Fig. 3a-c; Table S4).

**Figure 3:**
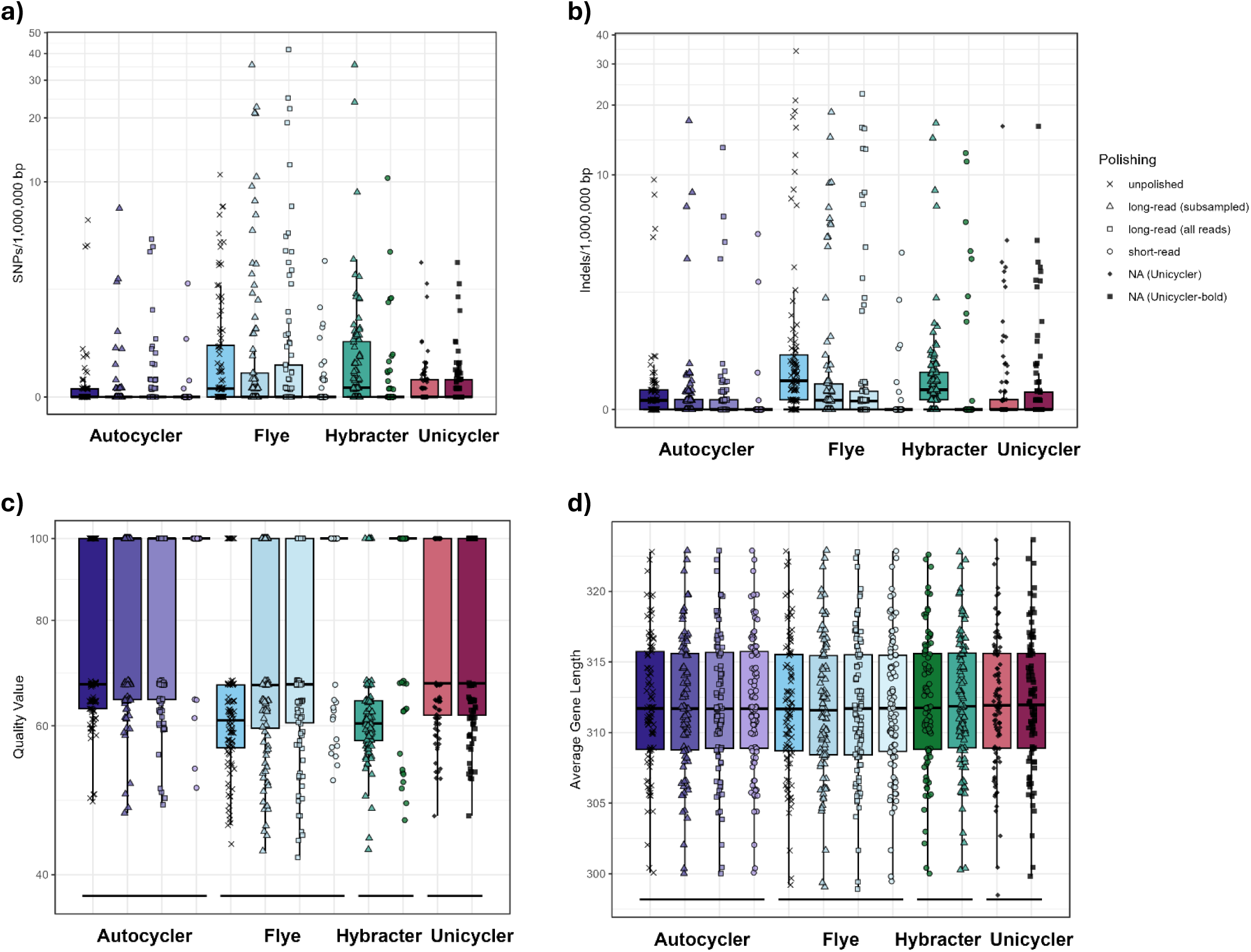
Assembly accuracy for different assembler/polisher combinations. **a)** Single nucleotide substitution errors (SNPs) and **b)** insertion/deletions (indels) identified by re-aligning Illumina short-reads, **c)** quality value as annotated by Freebayes(48) from Pypolca(17) and **d)** mean gene length from CheckM2(37) of 12 different assembler/polisher combinations. The y-axes in a), b) and c) are transformed using a pseudo-log scale to facilitate plotting zero values given log(0) is undefined.

#### Medaka long-read polishing offers small improvements in accuracy for long-read assemblies, although short-read polishing is still marginally more accurate

Medaka long-read polishing (with un-subsampled reads) improved accuracy for Autocycler and Flye by improving QV and reducing indels (from median Q67 to Q100 [Wilcoxon signed rank *p=0.007*], and Q61 to Q67 [*p<0.0001*], and 0.18 indels/Mb to 0 [*p=0.006*], and 0.57 indels/Mb to 0.17 [*p<0.001*], respectively), but there was no evidence of reducing SNPs (*p=1* for both Autocycler and Flye). There was some statistical evidence that Medaka long-read polishing using un-subsampled long-reads was marginally better at reducing indels for Autocycler assemblies than using subsampled reads (change vs Autocycler of median 0 indels/Mb [IQR: -0.19-0; range: -1.64-3.61] for un-subsampled reads, compared to a change of 0 [IQR: -0.18-0; range: -1.09-7.60] indels/Mb, Wilcoxon signed rank *p=0.019*; Fig.3; Table S3). However, this very small difference is not reflected in the medians/IQR of indels/Mb as most isolates had 0 indels (57% [52/92] for Autocycler + Medaka [subsampled] and 65% [60/92] for Autocycler + Medaka [un-subsampled]).

Short-read polished Autocycler assemblies were more accurate than the best long-read polished Autocycler assemblies (Autocycler + Medaka [un-subsampled]) (change vs unpolished Autocycler of median 0 [IQR: -0.16-0] SNPs/Mb, -0.18 [-0.39-0] indels/Mb, and Q32.6 (Q0-Q35.9) for short-read polishing vs median change 0 [0-0] SNPs/Mb, 0 [-0.19-0] indels/Mb, and Q0 (Q0-Q6.15) for Medaka (un-subsampled) polishing, pairwise Wilcoxon signed rank *p=0.0002*, *p<0.0001 and p<0.0001*, respectively; Fig 3; Table S4). However, the absolute difference was small, and affected only the worst-performing quartile of isolates. The majority, 55% (51/92), of Autocycler + Medaka (un-subsampled reads) polished assemblies had 0 errors (QV100), and only 4% (4/92) of genome sequences had >10 SNPs or indels in the entire assembly, compared with 95% (87/92) of short-read polished Autocycler assemblies having 0 errors and two genome sequences with >10 SNPs or indels (Figs. 3a-c; Table S4).

#### Mean gene length is slightly shorter for Flye assemblies, and is not corrected by long or short-read polishing

Mean gene length was assessed as a further measure of accuracy, as small errors can result in coding sequence truncation, and shorter average gene length. While there was some statistical evidence of a difference in mean gene length between different assembler/polisher combinations, with unpolished and long-read polished Flye assemblies having a slightly shorter mean gene length compared to other assembler (Friedman’s *p<0.0001*; all pairwise Wilcoxon signed rank *p<0.0001-p=0.01* compared to all other assemblers), the difference was small in magnitude (median of the mean gene length across all isolates of 312bp [IQR: 308-315bp] for Flye + Medaka (subsampled) polishing, vs 312bp [309-316bp] for all other non-Flye assemblers; Fig. 3d).

#### Gene annotation for MLST loci, resistance, virulence and stress genes is equivalent for long-read and hybrid assemblies

There was no evidence of a difference in the numbers of key resistance, virulence and stress genes identified by AMRFinder Plus in assemblies generated by any assembler/polisher combination (Friedman’s *p=0.209* for resistance, *p=0.736* for virulence, and *p=0.687* for stress genes; all pairwise Wilcoxon signed-rank *p=1*; Table S4). There was high concordance between assemblers on the presence/absence of specific gene variants (all pairwise McNemar’s *p>0.209*). There was also no evidence of a difference in the proportion of isolates with correctly assigned multi-locus sequence type (MLST; all pairwise McNemar’s *p=1*, Table S4). Hybracter (long; hybrid), Unicycler (normal; bold), and polished Flye assemblies were annotated with identical MLST-types for all 91 isolates belonging to a species with available MLST-typing schemes (i.e. all isolates except one *Serratia marcescens*). A single locus in one isolate was ‘uncertain’ for the unpolished Flye assembly ((*gapA*(∼2)), and another locus (*gyrB*(10)) was duplicated in a different isolate amongst Autocycler assemblies. Polishing did not correct this duplicated annotation, although the allele was correctly identified.

## Discussion

We evaluated three long-read only bacterial genome assemblers, three hybrid assemblers, and three polishers on 92 clinical Enterobacterales isolates. The consensus long-read assembler, Autocycler, produced the most structurally complete assemblies, circularising 95% of chromosomes. Plasmid reconstruction was comparable between all assemblers except Flye, which underperformed compared with other assemblers for most metrics. Autocycler with Medaka polishing was the most accurate long-read only assembler/polisher combination, with a median of 0 SNPs/indels compared to what we consider the ‘gold-standard’ hybrid assembly (i.e. short-read polished Autocycler assemblies). Long-read polishing of Autocycler and Flye assemblies offered small improvements in accuracy compared to unpolished assemblies, although short-read polishing still corrected marginally more errors. There was strong agreement in the annotation of seven-locus MLSTs, resistance, virulence and stress genes, and mean gene length across all assemblers.

It is not surprising that long-read assemblers circularise more chromosomes, as long-reads can resolve repetitive regions that short-reads may not. This explains why the long-read first hybrid assembler, Hybracter (hybrid), performed more similarly to other long-read assemblers than Unicycler, which uses short-reads first to reconstruct overall structure. The ability of Autocycler to circularise eight chromosomes where non-consensus assemblers failed supports the utility of this software(57). Combining 20 input assemblies in Autocycler may reduce the effects of stochastic variation in individual assemblers. The 2/92 isolates where Autocycler produced fragmented assemblies, while its some input assemblies were complete, are noteworthy. This result is perhaps attributable to regions of input assemblies that are too divergent to resolve, and highlights the need for an iterative approach, where a ‘fallback’ option is available in case of a highly fragmented Autocycler consensus assembly. This also emphasises the importance of quality controls (e.g.: checkM2) to flag highly fragmented assemblies, so that for these cases, manual curation of input assemblies, optimising parameters in the consensus process, or reversion to complete input assemblies may improve assembly.

Evaluation of chromosomal and plasmid sequence reconstruction is challenging due to the absence of a ‘ground truth’. For plasmids specifically, there is a risk of mislabelling plasmids by methods reliant on reference databases, which may be incomplete or contain misassembled plasmids. We therefore considered two reference plasmid sets generated from the study data. Compared with both reference sets, none of the six assemblers had ‘perfect’ concordance. Flye performed poorly compared to all other assemblers, missing or misassembling ∼45% of plasmids compared with 4-10% for other assemblers. Flye struggled particularly with small <10,000bp plasmids, as reported previously(16, 58). This emphasises the necessity of consensus methods like Autocycler(57), and separate plasmid recovery tools like Plassembler(31) to optimise plasmid reconstruction. The fact that Autocycler (including four Hybracter (long) input assemblies) reconstructed a slightly different set of plasmids to a single Hybracter (long/hybrid) assembly suggests complementarity between these methods, where Autocycler can overcome potential issues related to stochastic variation in individual assemblies. The replicon annotation differences between identical plasmids highlights the risks of relying on plasmid-annotation tools like MOB-suite for plasmid identification(59), and supports the use of network-based tools like PLING(60).

The small differences in nucleotide-level accuracy between long- and short-read polished Autocycler assemblies are likely not in coding regions that are key for downstream analyses. This is evidenced by the strong agreement in MLST profile, resistance, virulence and stress gene annotations, and mean gene length between assemblers.

The advantage of our study is that we consider a relatively large sample of real-world, clinically-relevant isolates. Specifically, our sample included predominantly *E. coli* and *K. pneumoniae*, which are the two most important Gram-negative species in England in terms of number of bloodstream infections and burden of AMR(61), and therefore our findings are relevant to public health surveillance in this setting. However, a trade-off with this is the absence of ‘ground truth’ sequences against which to evaluate our assemblies. Other limitations include the empirical assessment of nucleotide-level accuracy, through aligning short-reads to assemblies. Both SNPs and indels were still present in a small number of short-read polished assemblies, potentially representing a baseline level of errors in either Illumina reads or read mapping, and leading to possible overestimation of the error rate of long-read only assemblies. A further limitation is that the performance of Autocycler as a consensus method depends on its input assemblies. Twenty input assemblies were used here, requiring substantial computational time (13,428 CPUh), mostly due to generating assemblies, and resulted in a high carbon footprint, equivalent to driving 164 miles (see Environmental Impact Statement). Furthermore, a closed consensus chromosome was not achieved for 5% of isolates using default settings. Optimisation of Autocycler input assemblies and parameters, such as weighting contigs from certain ‘more reliable’ assembler, as done in more recent automated Autocycler v5 pipelines(20), could thus reduce computational load and improve performance. Incorporating a ‘fallback’ option in Autocycler pipelines, for example to revert to one of the complete input assemblies in cases of a highly fragmented Autocycler consensus, may also be of benefit. Finally, generalisability to other bacterial species is limited. Other species may be less-well represented than *E. coli* and *Klebsiella* spp. in the machine-learning training datasets for basecalling (Dorado) and polishing (Medaka) software, producing potentially different error rates.

## Conclusions

This assembly comparison is the first benchmarking study to demonstrate structural completeness and accuracy of Nanopore super-high accuracy long-read only bacterial genome assemblies on 92 clinical Enterobacterales isolates, compared with hybrid assembly. The automated consensus long-read assembler, Autocycler, accurately reconstructed assemblies, including plasmids, for these isolates, and is a promising tool for integrating Nanopore long-read only assemblies into an automatable computational pipeline for public health genomics. Ongoing innovation in Nanopore sequencing technology and bioinformatic software may enable further improvements and should continue to be evaluated by the bioinformatics community.

## Environmental Impact Statement

The Nextflow assembly pipeline used for this work ran in 72h on two AMD EPYC 9J14 96-Core Processors (188 total CPUs; 13,428 CPUh), and drew 124.46 kWh. Using Cloud infrastructure based in the United Kingdom, this had a carbon footprint of 28.76 kgCO2e, equivalent to 2.61 tree-years, or 164 km in a car (calculated using green-algorithms.org v3.0(62)). This is a lower bound estimate of the carbon footprint of this work, as it does not account for compute used in pipeline development, downstream statistical analyses, or the energy required to power display screens. The carbon footprint and wider environmental impact of sample processing shipping has also not been accounted for.

## Conflict of interest

The authors have no conflicts of interest to declare.

## Funding information

This study/research is supported/funded by the National Institute for Health Research (NIHR) Health Protection Research Unit in Healthcare Associated Infections and Antimicrobial Resistance (NIHR207397), a partnership between the UK Health Security Agency (UKHSA) and the University of Oxford. This work was also supported by the UKHSA and the NIHR Oxford Biomedical Research Centre (BRC) and the UKHSA PhD Funding Competition. The cloud compute infrastructure for this work was donated by Oracle Corporation Infrastructure. The views expressed are those of the authors and not necessarily those of the NIHR, UKHSA or the Department of Health and Social Care.

## Ethical approval and consent to participate

This work has been reviewed and approved by the UKHSA Research Ethics & Governance Group (reference NR0429).

## Consent for publication

All authors give consent for publication of this work. No further consent for publication was required as this work does not include patient identifiable information.

## Author contributions

NS, SL, SH, DC, ASW, JR, KLH, AL, DW, RH and CSB were involved in conceptualisation, funding acquisition, project administration, provision or resources and supervision. VP, GR, KH, CRJ and NEKSUS consortium members were involved in isolate collection and processing. Methodological development and validation of bioinformatic methods and software was done by DN under the supervision of SL and NS. DN, SL and NS were involved with data curation, analysis, investigation, visualisation and writing/editing. All authors approved the final draft

## Acknowledgments

The authors would also like to acknowledge all participating laboratories in the NEKSUS consortium who were responsible for isolate collection, Zeynab Yusuf from UKHSA for her role in sample transportation from UKHSA to Oxford, laboratory and bioinformatician colleagues at the Modernising Medical Microbiology Unit at the University of Oxford for support in methodological development and execution, as well as GENEWIZ Germany GmbH (Leipzig, Germany) for performing long- and short-read sequencing.

Individuals within the NEKSUS consortium group authorship are (listed alphabetically):

- Alan McNally (University Hospitals Birmingham NHS Foundation Trust)
- Caroline Cullerton (The Newcastle-upon-Tyne Hospitals NHS Foundation Trust)
- Gabriella Shanks (Barts Heath NHS Trust)
- James Price (University Hospital Sussex NHS Foundation Trust)
- Jasvir Nahl (Leeds Teaching Hospitals NHS Trust)
- Jenny Bradbury (UKHSA)
- Jonathan Lambourne (Barts Health NHS Trust)
- Julie Samuel (The Newcastle-upon-Tyne Hospitals NHS Foundation Trust)
- Jumoke Sule (UKHSA/ Cambridge University Hospitals NHS Foundation Trust)
- Ian Butler (Barts Health NHS Trust)
- Kavita Sethi (Leeds Teaching Hospitals NHS Trust)
- Mark Garvey (University Hospitals Birmingham NHS Foundation Trust)
- Martin Williams (University Hospitals Bristol and Weston NHS Foundation Trust)
- Nicholas Brown (Cambridge University Hospitals NHS Foundation Trust)
- Nicola Childs (North Bristol NHS Trust)
- Paul Randell (University Hospital Sussex NHS Foundation Trust)
- Poorvi Patel (Cambridge University Hospitals NHS Foundation Trust)
- Samuel Stafford (North Bristol NHS Trust)
- Samuel Tetley (University Hospital Sussex NHS Foundation Trust)
- Simon Eccles (Manchester University Hospitals NHS Foundation Trust)

## Supplementary Figures

**Supplementary Figure S1:**
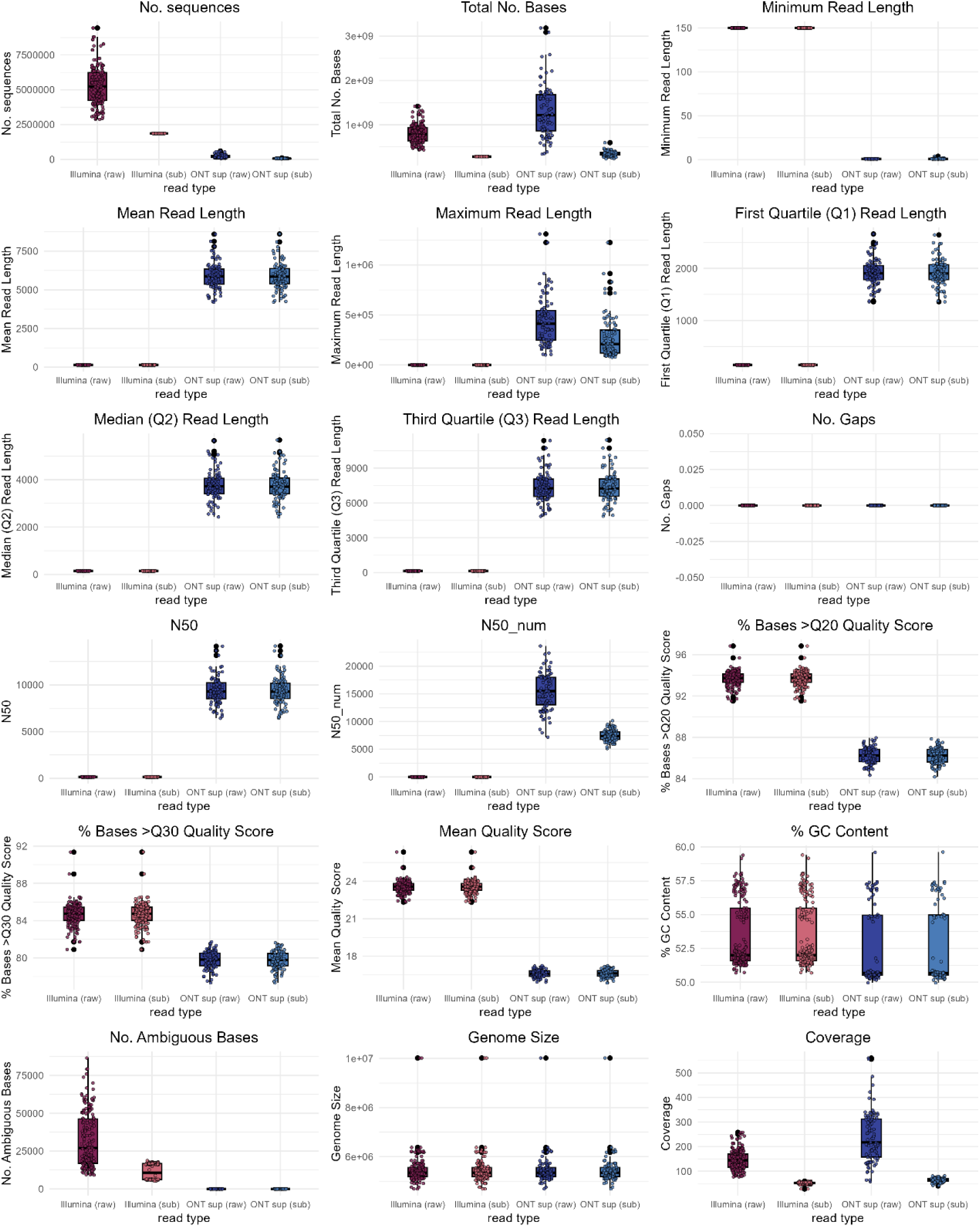
Quality control metrics of raw and subsampled Illumina short-reads and Dorado v5.0.0 super accurate basecalled Nanopore long-reads. Showing long-read subsampled set 1 (of 4) for the 92 pure culture isolates. N50 and N50_num (or L50) are both measures of sequence contiguity(63). N50 is the sequence length of the shortest contig at 50% of the total assembly length. N50_num is defined as the count of the smallest number of contigs whose added length makes up at least half of genome size.

**Supplementary Figure S2:**
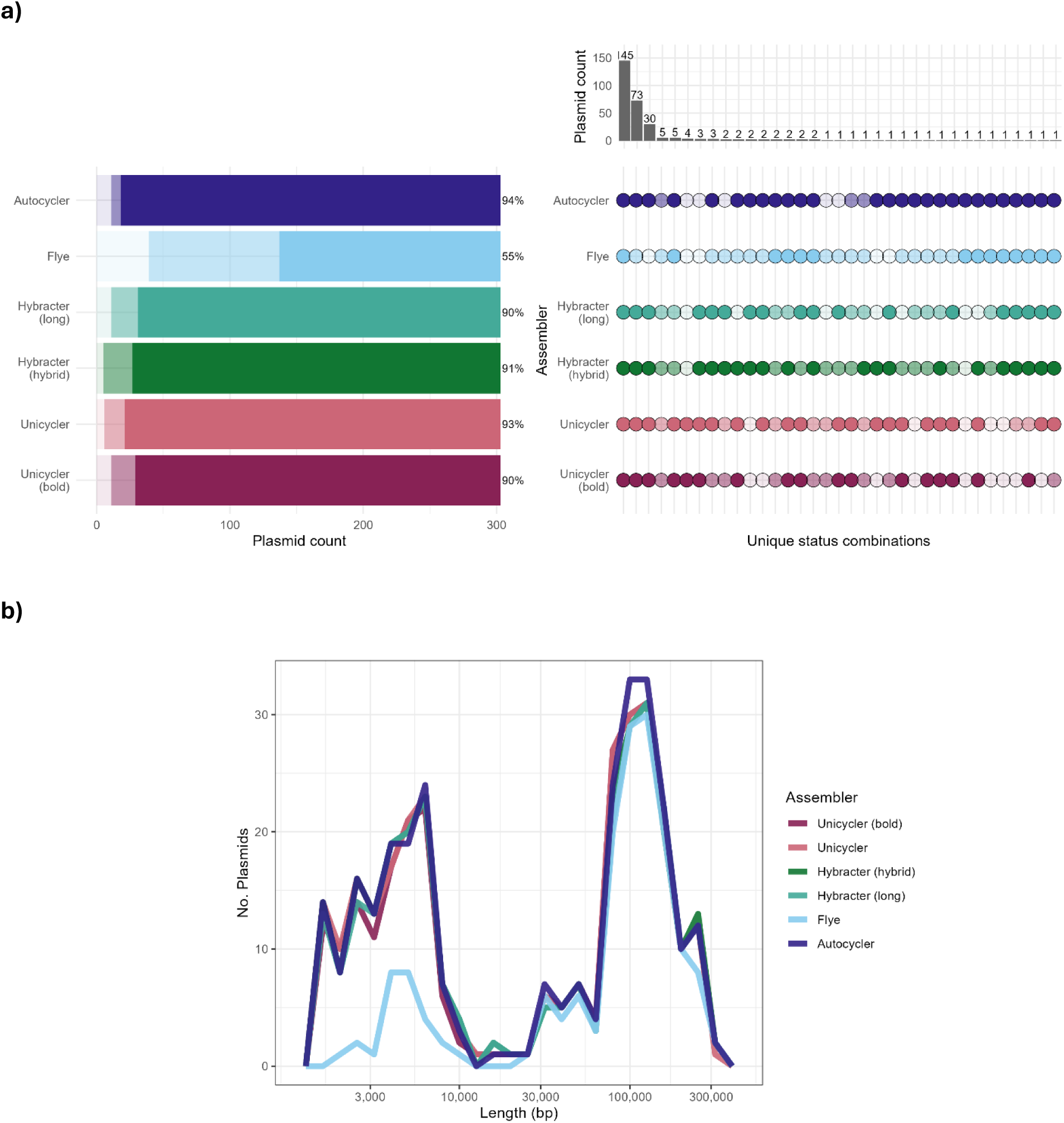
Plasmid sequence reconstruction for 92 Enterobacterales isolates by different long-read only and hybrid assemblers, using the manually-curated consensus ‘reference’ plasmid set (n=303 plasmids). Reference plasmids in the manually curated set are circular contigs between 1,000-400,000bp in length that are present in at least 2 assemblers with a matching length (±10%) and mash distance (<0.025). **a)** Upset plot showing assembly status combinations of plasmids across assemblers. Dark circles/bars indicate ‘present’ plasmids where length (±10%), mash distance (<0.025) and circularity all matched the ‘reference’ plasmid, lighter colours indicate misassembled plasmids, where the length difference was >10%, mash distance >0.025, or the contig was non-circular and the palest shades indicate absent plasmids, where no contig was found matching other plasmids in the reference plasmid set. **b)** Frequency polygon of length distribution of ‘present’ plasmids by assembler.

**Supplementary Figure S3:**
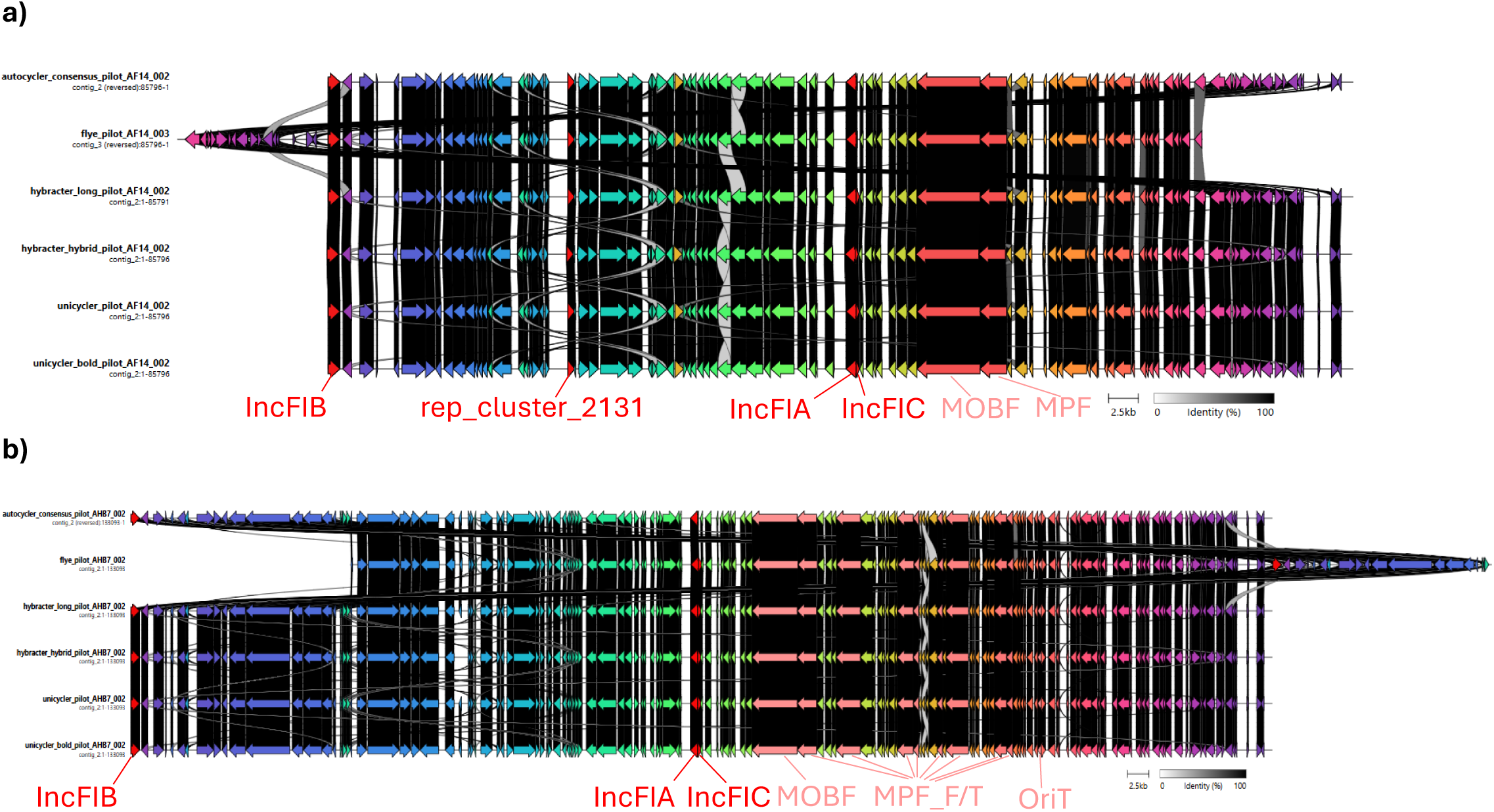
Clinker plots of highly similar plasmids with different MOB-suite annotations. Replicon annotations are shown in bright red and labelled. Other mobility- and replication-associated plasmid machinery are shown in pale red and labelled. **a)** An 85,796bp IncFIA, IncFIB, IncFIC, rep_cluster_2131 plasmid sequence (isolate AF14) with a missing IncFIC annotation in the Autocycler and Flye assemblies (top 2), despite a mash distance of 0 between Autocycler and Hybracter (hybrid) assemblies. **b)** A 133,309bp IncFIA, IncFIB, IncFIC plasmid sequence (isolate AHB7) with the IncFIC replicon annotation missing from the Autocycler plasmid sequence, despite a mash distance of 0 between the Autocycler and Hybracter (hybrid) plasmid sequences. Note the Autocycler plasmid sequence is reversed and the Flye plasmid has a different starting point for both plasmids. The Flye plasmid is also reversed in a) compared to the bottom 4 assemblers’ plasmids.

**Supplementary Table S1:** Species of the 92 pure culture Enterobacterales isolates, as assigned by Kraken2. (**40**).

| Species | Count (percentage) |
| --- | --- |
| *Escherichia coli* | 58 (63%) |
| *Klebsiella pneumoniae* | 21 (23%) |
| *Klebsiella oxytoca* | 6 (7%) |
| *Klebsiella aerogenes* | 2 (2%) |
| *Enterobacter hormaechei* | 2 (2%) |
| *Citrobacter freundii* | 1 (1%) |
| *Citrobacter portucalensis* | 1 (1%) |
| *Serratia marcescens* | 1 (1%) |

**Supplementary Table S2:**
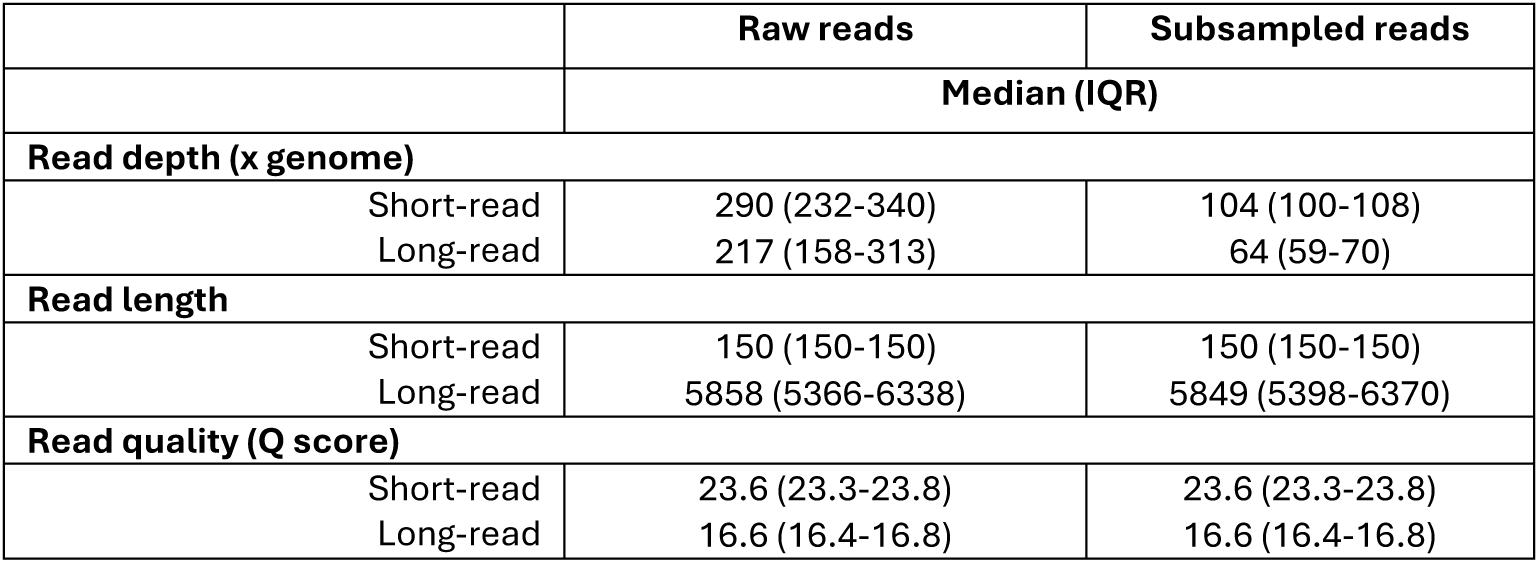
Raw and subsampled sequencing read metrics for Illumina short-read and Nanopore long-read sequences for 92 pure culture Enterobacterales isolates.

**Supplementary Table S3:** Plasmid reconstruction accuracy of different long-read only and hybrid assemblers for Dorado v5.0.0 super accurate basecalled Nanopore long-reads. Plasmid reconstruction is compared to a manually-curated reference set of ‘consensus’ plasmids (n=303), where ‘consensus’ plasmids were circular contigs 1,000-400,000bp in length present across at least 2 assemblers with a similar length (±10%) and close mash distance (<0.025).

|  | Assembler |  |  |  |  |  | p-value† |
| --- | --- | --- | --- | --- | --- | --- | --- |
|  | Autocycler<br>n (%) | Flye<br>n (%) | Hybracter (long)<br>n (%) | Hybracter (hybrid)<br>n (%) | Unicycler<br>n (%) | Unicycler (bold)<br>n (%) |  |
| **Present\* plasmids** | 285 (94.1%) | 166 (54.8%) | 272 (89.8%) | 276 (91.1%) | 282 (93.1%) | 274 (90.4%) | <0.0001 |
| **Misassembled\*\* plasmids** |  |  |  |  |  |  |  |
| Non-circular | 0 (0%) | 18 (5.9%) | 12 (4.0%) | 13 (4.3%) | 5 (1.7%) | 3 (1%) |  |
| Length mismatch | 7 (2.3%) | 50 (16.5%) | 1 (0.3%) | 2 (0.7%) | 6 (2.0%) | 7 (2.3%) |  |
| Non-circular and length mismatch | 0 (0%) | 30 (9.9%) | 7 (2.3%) | 7 (2.3%) | 4 (1.3%) | 8 (2.6%) |  |
| **Absent plasmids** | 11 (3.6%) | 39 (12.9%) | 11 (3.6%) | 5 (1.7%) | 6 (2.0%) | 11 (3.6%) |  |

\*‘Present’ plasmids are defined as contigs 1,000-400,000bp in length meeting all three match criteria: circular, length ( $\pm 10\%$ ) and mash distance ( $<0.025$ ) of a the manually curated reference set of plasmids.
\*\*Misassembled plasmids are defined as contigs that failed to meet at least 1 of the matching criteria, but could still be matched to the reference set based on a more distant mash distance.
\*\*\*Absent plasmids were cases where only the circularity matched, or where, for an assembler, no contig could be matched to the rest of the reference plasmids match set based on mash distance.
†p-value for Fleiss’ Kappa test for uneven proportions of ‘present’ plasmids across all assemblers.

**Supplementary Table S4:**
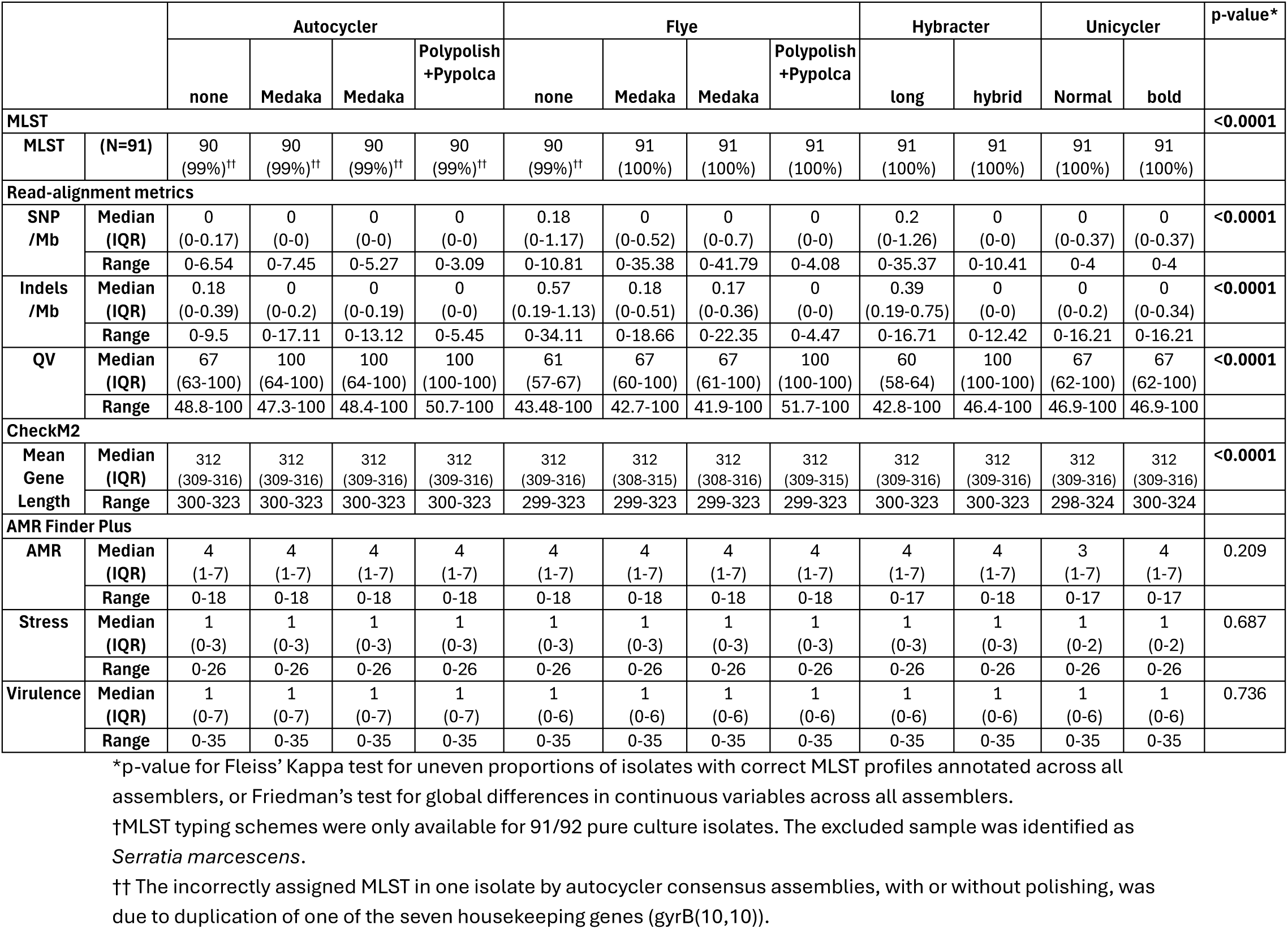
Nucleotide-level accuracy of 12 assembler-polisher combinations (7 long-read only, 5 hybrid). Read-alignment metrics were derived by aligning Illumina short-reads to each assembler-polisher combination and variant calling with Feebayes(48) from Pypolca. Mean gene length is derived from CheckM2(37) output files. 7-locus MLST is annotated by mlst(39), and key resistance, virulence and stress genes by AMRFinder Plus(64).

## Notes

https://figshare.com/account/home#/projects/253775

https://github.com/oxfordmmm/NEKSUS_ont_hybrid_assembly_comparison

### Competing Interest Statement

The authors have declared no competing interest.

## References

1. Sanderson ND, Kapel N, Rodger G, Webster H, Lipworth S, Street TL, et al. Comparison of R9.4.1/Kit10 and R10/Kit12 Oxford Nanopore flowcells and chemistries in bacterial genome reconstruction. Microb Genom. 2023;9(1). 10.1099/mgen.0.000910

2. Hall MB, Wick RR, Judd LM, Nguyen AN, Steinig EJ, Xie O, et al. Benchmarking reveals superiority of deep learning variant callers on bacterial nanopore sequence data. Elife. 2024;13. 10.7554/eLife.98300

3. Sereika M, Kirkegaard RH, Karst SM, Michaelsen TY, Sørensen EA, Wollenberg RD, et al. Oxford Nanopore R10.4 long-read sequencing enables the generation of near-finished bacterial genomes from pure cultures and metagenomes without short-read or reference polishing. Nat Methods. 2022;19(7):823–6. 10.1038/s41592-022-01539-7

4. Ni Y, Liu X, Simeneh ZM, Yang M, Li R. Benchmarking of Nanopore R10.4 and R9.4.1 flow cells in single-cell whole-genome amplification and whole-genome shotgun sequencing. Comput Struct Biotechnol J. 2023;21:2352–64. 10.1016/j.csbj.2023.03.038

5. Foster-Nyarko E, Cottingham H, Wick RR, Judd LM, Lam MMC, Wyres KL, et al. Nanopore-only assemblies for genomic surveillance of the global priority drug-resistant pathogen, Klebsiella pneumoniae. Microb Genom. 2023;9(2). 10.1099/mgen.0.000936

6. Wick RR, Judd LM, Holt KE. Assembling the perfect bacterial genome using Oxford Nanopore and Illumina sequencing. PLoS Comput Biol. 2023;19(3):e1010905. 10.1371/journal.pcbi.1010905

7. Wick RR, Judd LM, Gorrie CL, Holt KE. Unicycler: Resolving bacterial genome assemblies from short and long sequencing reads. PLOS Computational Biology. 2017;13(6):e1005595. 10.1371/journal.pcbi.1005595

8. Heather JM, Chain B. The sequence of sequencers: The history of sequencing DNA. Genomics. 2016;107(1):1–8. 10.1016/j.ygeno.2015.11.003

9. Wang Y, Yang Q, Wang Z. The evolution of nanopore sequencing. Front Genet. 2014;5:449. 10.3389/fgene.2014.00449

10. Simar SR, Hanson BM, Arias CA. Techniques in bacterial strain typing: past, present, and future. Curr Opin Infect Dis. 2021;34(4):339–45. 10.1097/qco.0000000000000743

11. Castaneda-Barba S, Top EM, Stalder T. Plasmids, a molecular cornerstone of antimicrobial resistance in the One Health era. Nat Rev Microbiol. 2024;22(1):18–32. 10.1038/s41579-023-00926-x

12. Dimitriu T. Evolution of horizontal transmission in antimicrobial resistance plasmids. Microbiology (Reading). 2022;168(7). 10.1099/mic.0.001214

13. Khezri A, Avershina E, Ahmad R. Hybrid Assembly Provides Improved Resolution of Plasmids, Antimicrobial Resistance Genes, and Virulence Factors in Escherichia coli and Klebsiella pneumoniae Clinical Isolates. Microorganisms. 2021;9(12). 10.3390/microorganisms9122560

14. Arredondo-Alonso S, Willems RJ, van Schaik W, Schurch AC. On the (im)possibility of reconstructing plasmids from whole-genome short-read sequencing data. Microb Genom. 2017;3(10):e000128. 10.1099/mgen.0.000128

15. Sanderson ND, Hopkins KMV, Colpus M, Parker M, Lipworth S, Crook D, et al. Evaluation of the accuracy of bacterial genome reconstruction with Oxford Nanopore R10.4.1 long-read-only sequencing. Microb Genom. 2024;10(5). 10.1099/mgen.0.001246

16. Abdel-Glil MY, Brandt C, Pletz MW, Neubauer H, Sprague LD. High intra-laboratory reproducibility of nanopore sequencing in bacterial species underscores advances in its accuracy. Microbial Genomics. 2025;11(3).10.1099/mgen.0.001372

17. Bouras G, Judd LM, Edwards RA, Vreugde S, Stinear TP, Wick RR. How low can you go? Short-read polishing of Oxford Nanopore bacterial genome assemblies. Microb Genom. 2024;10(6). 10.1099/mgen.0.001254

18. De Maio N, Shaw LP, Hubbard A, George S, Sanderson ND, Swann J, et al. Comparison of long-read sequencing technologies in the hybrid assembly of complex bacterial genomes. Microb Genom. 2019;5(9). 10.1099/mgen.0.000294

19. Bouras G, Houtak G, Wick RR, Mallawaarachchi V, Roach MJ, Papudeshi B, et al. Hybracter: enabling scalable, automated, complete and accurate bacterial genome assemblies. Microb Genom. 2024;10(5). 10.1099/mgen.0.001244

20. Wick RR. Autocycler. 2025.

21. Wick RR, Judd LM, Cerdeira LT, Hawkey J, Méric G, Vezina B, et al. Trycycler: consensus long-read assemblies for bacterial genomes. Genome Biology. 2021;22(1):266. 10.1186/s13059-021-02483-z

22. Zhou A, Lin T, Xing J. Evaluating nanopore sequencing data processing pipelines for structural variation identification. Genome Biology. 2019;20(1):237. 10.1186/s13059-019-1858-1

23. illumina. bcl2fastq2 Conversion Software v2.20. 2017.

24. Oxford Nanopore Technologies. Dorado v0.9 2024 [Available from: https://github.com/nanoporetech/dorado?tab=readme-ov-file#alignment.

25. Shen W, Le S, Li Y, Hu F. SeqKit: A Cross-Platform and Ultrafast Toolkit for FASTA/Q File Manipulation. PLOS ONE. 2016;11(10):e0163962. 10.1371/journal.pone.0163962

26. Hall MB. Rasusa: Randomly subsample sequencing reads to a specified coverage. Journal of Open Source Software. 2022; 7(69):3941.10.21105/joss.03941

27. Kolmogorov M, Yuan J, Lin Y, Pevzner P. Assembly of Long Error-Prone Reads Using Repeat Graphs. Nature Biotechnology. 2019.doi:10.1038/s41587-019-0072-8

28. Koren S, Walenz BP, Berlin K, Miller JR, Bergman NH, Phillippy AM. Canu: scalable and accurate long-read assembly via adaptive k-mer weighting and repeat separation. Genome Res. 2017;27(5):722–36. 10.1101/gr.215087.116

29. Vaser R, Šikić M. Time- and memory-efficient genome assembly with Raven. Nature Computational Science. 2021;1(5):332–6. 10.1038/s43588-021-00073-4

30. Li H. Minimap and miniasm: fast mapping and de novo assembly for noisy long sequences. Bioinformatics. 2016;32(14):2103–10. 10.1093/bioinformatics/btw152

31. Bouras G, Sheppard AE, Mallawaarachchi V, Vreugde S. Plassembler: an automated bacterial plasmid assembly tool. Bioinformatics. 2023;39(7). 10.1093/bioinformatics/btad409

32. Lee JY, Kong M, Oh J, Lim J, Chung SH, Kim JM, et al. Comparative evaluation of Nanopore polishing tools for microbial genome assembly and polishing strategies for downstream analysis. Sci Rep. 2021;11(1):20740. 10.1038/s41598-021-00178-w

33. Wick RR, Holt KE. Polypolish: Short-read polishing of long-read bacterial genome assemblies. PLOS Computational Biology. 2022;18(1):e1009802. 10.1371/journal.pcbi.1009802

34. Zimin AV, Salzberg SL. The genome polishing tool POLCA makes fast and accurate corrections in genome assemblies. PLOS Computational Biology. 2020;16(6):e1007981. 10.1371/journal.pcbi.1007981

35. Chklovski. CheckM2. 1.1.0 ed 2025.

36. Parks DH, Imelfort M, Skennerton CT, Hugenholtz P, Tyson GW. CheckM: assessing the quality of microbial genomes recovered from isolates, single cells, and metagenomes. Genome Res. 2015;25(7):1043–55. 10.1101/gr.186072.114

37. Chklovski A, Parks DH, Woodcroft BJ, Tyson GW. CheckM2: a rapid, scalable and accurate tool for assessing microbial genome quality using machine learning. Nature Methods. 2023;20(8):1203–12. 10.1038/s41592-023-01940-w

38. Schwengers O, Jelonek L, Dieckmann MA, Beyvers S, Blom J, Goesmann A. Bakta: rapid and standardized annotation of bacterial genomes via alignment-free sequence identification. Microbial Genomics. 2021;7(11).10.1099/mgen.0.000685

39. Seemann, Torsten. mlst. 2.23.0 ed: Github.

40. Wood DE, Lu J, Langmead B. Improved metagenomic analysis with Kraken 2. Genome Biology. 2019;20(1):257. 10.1186/s13059-019-1891-0

41. Robertson J, Nash JHE. MOB-suite: software tools for clustering, reconstruction and typing of plasmids from draft assemblies. Microb Genom. 2018;4(8). 10.1099/mgen.0.000206

42. Robertson J, Bessonov K, Schonfeld J, Nash JHE. Universal whole-sequence-based plasmid typing and its utility to prediction of host range and epidemiological surveillance. Microb Genom. 2020;6(10). 10.1099/mgen.0.000435

43. Ondov BD, Starrett GJ, Sappington A, Kostic A, Koren S, Buck CB, et al. Mash Screen: high-throughput sequence containment estimation for genome discovery. Genome Biology. 2019;20(1):232. 10.1186/s13059-019-1841-x

44. Ondov BD, Treangen TJ, Melsted P, Mallonee AB, Bergman NH, Koren S, et al. Mash: fast genome and metagenome distance estimation using MinHash. Genome Biology. 2016;17(1):132. 10.1186/s13059-016-0997-x

45. Csárdi G, Nepusz T, Traag V, Horvát S, Zanini F, Noom D, et al. igraph: Network Analysis and Visualization in R. R package version 2.1.4 ed 2025.

46. Csardi G, Nepusz T. The igraph software package for complex network research. InterJournal, Complex Systems. 2006;1695

47. Li H. Aligning sequence reads, clone sequences and assembly contigs with BWA-MEM. arXiv. 2013:1303.3997

48. Garrison E, Marth G. Haplotype-based variant detection from short-read sequencing. arXiv. 2012:e1207.3907

49. R Core Team. R: A Language and Environment for Statistical Computing. 4.4.1 ed 2021.

50. Wickham H. ggplot2: Elegant Graphics for Data Analysis. Verlag New York: Springer; 2016.

51. Hadley Wickham, Averick M, Bryan J, Chang W, McGowan LDA, François R, et al. Welcome to the tidyverse. Journal of Open Source Software. 2019;4(43):1686. 10.21105/joss.01686

52. Auguie B, Antonov A. gridExtra: Miscellaneous Functions for “Grid” Graphics 2.3 ed 2017.

53. Wilke CO. cowplot: Streamlined Plot Theme and Plot Annotations for ‘ggplot2’. 2024.

54. Revelle W. psych: Procedures for Psychological, Psychometric, and Personality Research R package version 2.5.6 ed. Evanston, Illinois: Northwestern University; 2025.

55. Gamer M, Lemon J, Fellows I, Singh P. irr: Various Coefficients of Interrater Reliability and Agreement. 0.84.1 ed 2019.

56. Gilchrist CLM, Chooi Y-H. clinker & clustermap.js: automatic generation of gene cluster comparison figures. Bioinformatics. 2021;37(16):2473–5. 10.1093/bioinformatics/btab007

57. Wick RR, Howden BP, Stinear TP. Autocycler: long-read consensus assembly for bacterial genomes. bioRxiv. 2025. 10.1101/2025.05.12.653612

58. Wick RR, Judd LM, Wyres KL, Holt KE. Recovery of small plasmid sequences via Oxford Nanopore sequencing. Microb Genom. 2021;7(8). 10.1099/mgen.0.000631

59. Douarre PE, Mallet L, Radomski N, Felten A, Mistou MY. Analysis of COMPASS, a New Comprehensive Plasmid Database Revealed Prevalence of Multireplicon and Extensive Diversity of IncF Plasmids. Front Microbiol. 2020;11:483. 10.3389/fmicb.2020.00483

60. Frolova D, Lima L, Roberts L, Bohnenkämper L, Wittler R, Stoye J, et al. Applying rearrangement distances to enable plasmid epidemiology with pling. bioRxiv. 2024:2024.06.12.598623. 10.1101/2024.06.12.598623

61. UK Health Security Agency. English surveillance programme for antimicrobial utilisation and resistance (ESPAUR) Report 2023 to 2024. 2024.

62. Lannelongue L, Grealey J, Inouye M. Green Algorithms: Quantifying the Carbon Footprint of Computation. Adv Sci (Weinh). 2021;8(12):2100707. 10.1002/advs.202100707

63. Wikipedia. N50, L50, and related statistics 2024 [Available from: https://en.wikipedia.org/wiki/N50,_L50,_and_related_statistics.

64. Feldgarden M, Brover V, Gonzalez-Escalona N, Frye JG, Haendiges J, Haft DH, et al. AMRFinderPlus and the Reference Gene Catalog facilitate examination of the genomic links among antimicrobial resistance, stress response, and virulence. Sci Rep. 2021;11(1):12728. 10.1038/s41598-021-91456-0

